# Bayesian adjustment for trend of colorectal cancer incidence in misclassified registering across Iranian provinces

**DOI:** 10.1101/340562

**Authors:** Sajad Shojaee, Nastaran Hajizadeh, Hadis Najafimehr, Luca Busani, Mohamad Amin Pourhoseingholi, Ahmad Reza Baghestani, Maryam Nasserinejad, Sara Ashtari, Mohammad Reza Zali

## Abstract

One of the problems in cancer registry of developing countries is misclassification error. This error leads to overestimation and underestimation of cancer rate in different provinces. The aim of this study is to use Bayesian method to correct for misclassification in registering cancer incidence in neighboring provinces of Iran. Incidence data of colorectal cancer were extracted from Iranian annual of national cancer registration reports 2005 to 2008 And Eighteen of the thirty Iranian provinces were selected to enter the Bayesian model and to correct their misclassification. Always a province with appropriate medical facilities is comparable to its neighbor or neighbors. Between years of 2005 and 2008, on the average, 28% misclassification was estimated between the province of East Azarbaijan and West Azarbayjan, 56% between the province of Fars and Hormozgan, 43% between the province of Isfahan and Charmahal and Bakhtyari, 46% between the province of Isfahan and Lorestan, 58% between the province of Razavi Khorasan and North Khorasan, 50% between the province of Razavi Khorasan and South Khorasan, 74% between the province of Razavi Khorasan and Sistan and Balochestan, 43% between the province of Mazandaran and Golestan, 37% between the province of Tehran and Qazvin, 45% between the province of Tehran and Markazi, 42% between the province of Tehran and Qom, 47% between the province of Tehran and Zanjan. Correcting the regional misclassification and obtaining the correct rates of cancer incidence in different regions is necessary for making cancer control and prevention programs and in healthcare resource allocation.

## Introduction

Colorectal cancer (CRC) is the third most common cancer among men (10.0% of the total) and the second in women (9.2% of the total) worldwide. Mortality is lower (694,000 deaths, 8.5% of the total) with more deaths (52%) in the less developed regions of the world, reflecting a poorer survival in these regions [1]. In Iran, CRC is the fourth most common type of cancer (the third most common cancer among females and the fifth among males), which accounts for 8.4% of total cancers in the country [2,3].

There is wide geographical variation in incidence across the world; the highest estimated rates being in Australia/New Zealand, and the lowest in Western Africa. Almost 55% of the cases occur in more developed regions. Of course, it is partly because of their advanced diagnostic and registration capabilities [1].

Inflammatory bowel disease, family history of CRC, obesity, dietary habits, smoking, physical inactivity [2,4], and diabetes [5] are well-known risk factors for CRC. Furthermore, environmental risk factors are found to play the most important role in the incidence and development of CRC [4]. So, people living in the same or adjacent areas which are imposed with the same environmental risk factors are expected to have similar cancer incidence rates.

Since cancer is a leading cause of morbidity and mortality worldwide [6,7], population-based and accurate information on its occurrence is extremely valuable as the foundation for identifying risk factors and making purposeful cancer prevention policies [8]. Cancer registries which are known as the main source of epidemiological data, which collects information regarding burden of cancers by recording the incidence, prevalence, survival and mortality of different cancers in a systematic manner [9–11]. Nowadays, their role has expanded into detecting the impact of interventions for cancer control, planning and evaluation of cancer screening programs, and specifying future needs for materials and manpower resources. But the existence of deficiencies in registering individuals information included patient’s permanent residence, primary site of tumor, date of diagnosis, and date of death [8], makes the registered data inaccurate to use in future planning.

In many developing countries such as Iran, health facilities are not distributed evenly throughout the country. So most cancer patients throughout the country prefer to get diagnostic and medical treatment services in capital of the country or in their neighboring facilitate provinces [12]. Some of them don’t mention their permanent residence and are registered in those provinces. It leads to misclassification error in cancer registry data. Misclassification error is the disagreement between the observed value and the true value in categorical data. As the evidence of existence of misclassification error in registering cancer incidence, the expected coverage of new cancer cases in different provinces can be mentioned; that the observed number of incidence is more than expected number in some provinces, and on the other hand, it is less than expected in a neighboring province [13]. It occurs while it is expected that the rate of cancer incidence be about the same in adjacent provinces; since people are adopt very similar lifestyle and traditions and are exposed to same environmental conditions.

There are two approaches to correct for misclassification error; the first approach is validating a small sample of data with rechecking medical records and extending the results to the target population [14]. The second approach is implementing Bayesian method. Bayesian method is a statistical approach that lets us to take our prior evidence into account in the analysis [15] with determining prior information for some of the parameters [16–18].

The aim of this study is to investigate the trend of colorectal cancer provinces of Iran after estimating the misclassification rate in registering cancer incidence by using Bayesian method and re-estimating the incidence rate in each province of Iran.

## Material and methods

Registering of cancer reports is obtainable from different references such as pathologies, hospitals, death certificates and etc. National registration programming of cancer cases from Iranian annual of national cancer registration report is extracted during 2005 to 2008 with software which was created by health ministry, until cancer cases are collected, registered and centralized for the past couple of years and is used for data analyses. Hence all new diagnosed cancer cases in temporary information bank are sent from medical universities to ministry of health periodically. Ministry of health after process of duplicating and coding the recorded cancers based on 10th revision of international coding of disease, this information is registered in permanent information bank. And all changes are sent to medical universities on specific duration until permanent information bank of medical universities is equalized with permanent information bank of health ministry. So each medical university has an observed number of cancer cases and also has an expected coverage of cancer cases that are considered to be 100 per 100000 except 2008 that was 113 per 100000. By dividing the observed number to the expected number of cancer cases, the percent of expected coverage for each province is calculated [20].

Since comparison of simple crude rate i.e. comparison of all cancer cases could make false images in total population regardless of age groups, age standardized rates (ASR) is calculated for all provinces of Iran using direct standardization method. The direct method for all provinces of Iran is based on, first selecting a criterion for the population and then calculating the desired outcome rate of this population using age specified rates at each of the two societies. At first, age groups were considered at level of 5 years. World standard population is the most common used standard population (*W_i_*). By dividing number of incident cases to person-years of observations, ASR is calculated per 100000 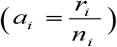. Finally for 4 age groups(0-14 years, 15-49 years, 50-69 years and over than70 years old) and for both genders, ASR is calculated in order to compare statistics on cancer internationally 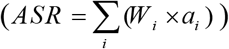 [20–22].

For entering the data to the Bayesian model two vectors *Y*_1_ and *Y*_2_ were used. Vector for the province, that, has an expected coverage less than 100% with exact ASR and vector for a neighboring province with a more than 100% expected coverage with ASR from the first group incorrectly labeled as being in the misclassified group. Subscript r is the number of covariate patterns for age and sex group combinations. A Poisson distribution was considered for count data and [19,22–24]. *Y*_1_ *Y*_2_

*Y*_1_ ≈ *Poisson*(*P_i_ μ*_*i*1_) and *Y*_2_ ≈ *Poisson*(*P_i_ μ*_*i*1_) the joint distribution of the count data *Y*_1_ and *Y*_2_ is proportional to:

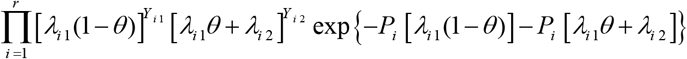

An informative beta prior distribution was assumed for *θ* as the probability of a data from the first group incorrectly registered in the misclassified group; so *θ* ~ *Beta*(*a, b*). For selecting prior value for the parameters of beta distribution, the calculated expected coverage for the medical university which has a less than 100% expected coverage was used as *b* and *a* was calculated with subtracting *b* from 100. Thus *a*/(*a* + *b*) which is the expectation of beta distribution converges to the misclassified rate. Variable U with binomial distribution, i.e. *U_i_* | *Y*_1_, *Y*_2_, *θ, λ*_1_, *λ*_2_ ~ *Binomial*(*Y*_*i*2_, *P_i_*) that 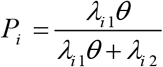 was considered as the number of events from the first group that are incorrectly registered in the misclassified group. Now if theta, *Y*_1_, *Y*_2_ to be unknown; we have:

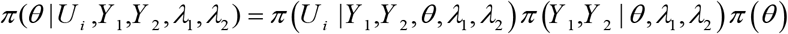

But since *Y*_1_, *Y*_2_ have known values of ASR on two neighboring provinces, then just theta is unknown and with employing a latent variable approach to correct the misclassification effect according to Paulino et al. [26,27], Liu et al. [28] and Stamey et al. [19] using a Gibbs sampling algorithm, the posterior appears in the following form:

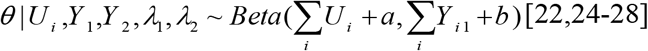

After estimating the misclassification rate between each two neighboring provinces, the rates of colorectal cancer incidence for each province were re-estimated and the trend of colorectal cancer were carried out during 2005 to 2008. All analyses were performed using R software version 3.3.1.

## Results

Registered cases of colorectal cancer have been included in the study for all provinces in Iran from 2005 to 2008. ASR of CRC incidence for men was 8.02 per 100,000 population (2255 cases) in 2005, whereas that year for women 7.4 per 100,000 (1801 cases). In over time, ASR of CRC incidence for men reached 12.7 per 100,000 population (3527 cases) in 2008 and for women to 11.12 per 100,000 (2658 cases) in the same year. The trend of CRC from 2005 to 2008 for both sexes is shown in Fig 1.

**Fig 1.**
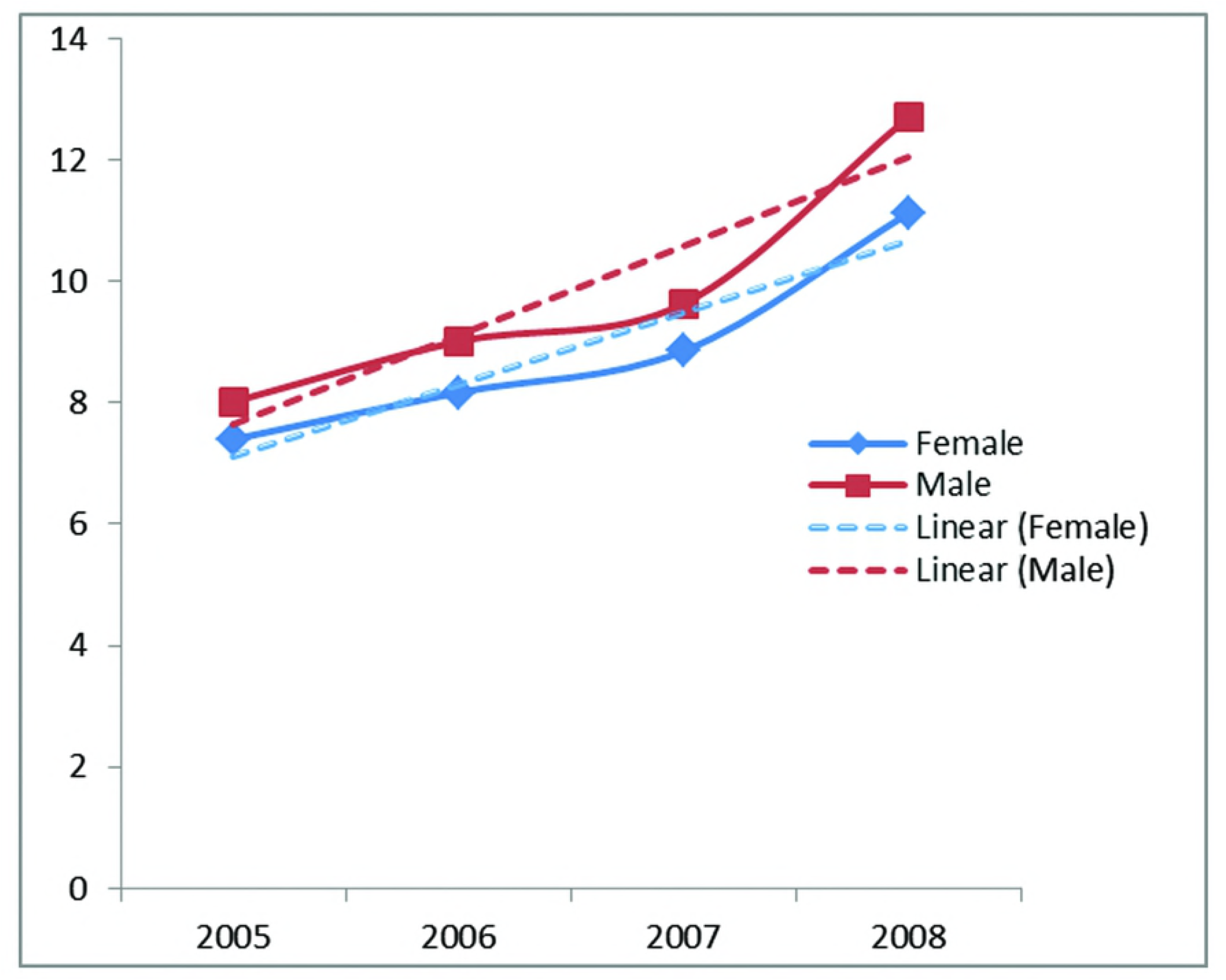
Age standardized rate of colorectal cancer incidence and its trend for male and female in Iran (2005-2008).

Among 30 provinces of Iran, 18 provinces where the number of cancer cases varied from their expected number were selected to correct the misclassification error in the register of CRC incidence in the neighbor provinces, based on the percentage of expected cancer coverage.

For example, the reported percentage of CRC expected coverage for Fars province as a province with suitable medical facilities and services was 120.8% in 2008. it means that Fars province have covered 20.8% of the new cases more than expected, while Hormozgan, which is adjacent to Fars, has a 19% expected coverage of cancer incidence, indicating clear misclassification in registering cancer cases. The expected coverage for all provinces in Iran between 2005 and 2008 is reported in Table 1. Also the estimated misclassification rate for all provinces in 2005-2008 is reported in Table 2.

**Table 1.**
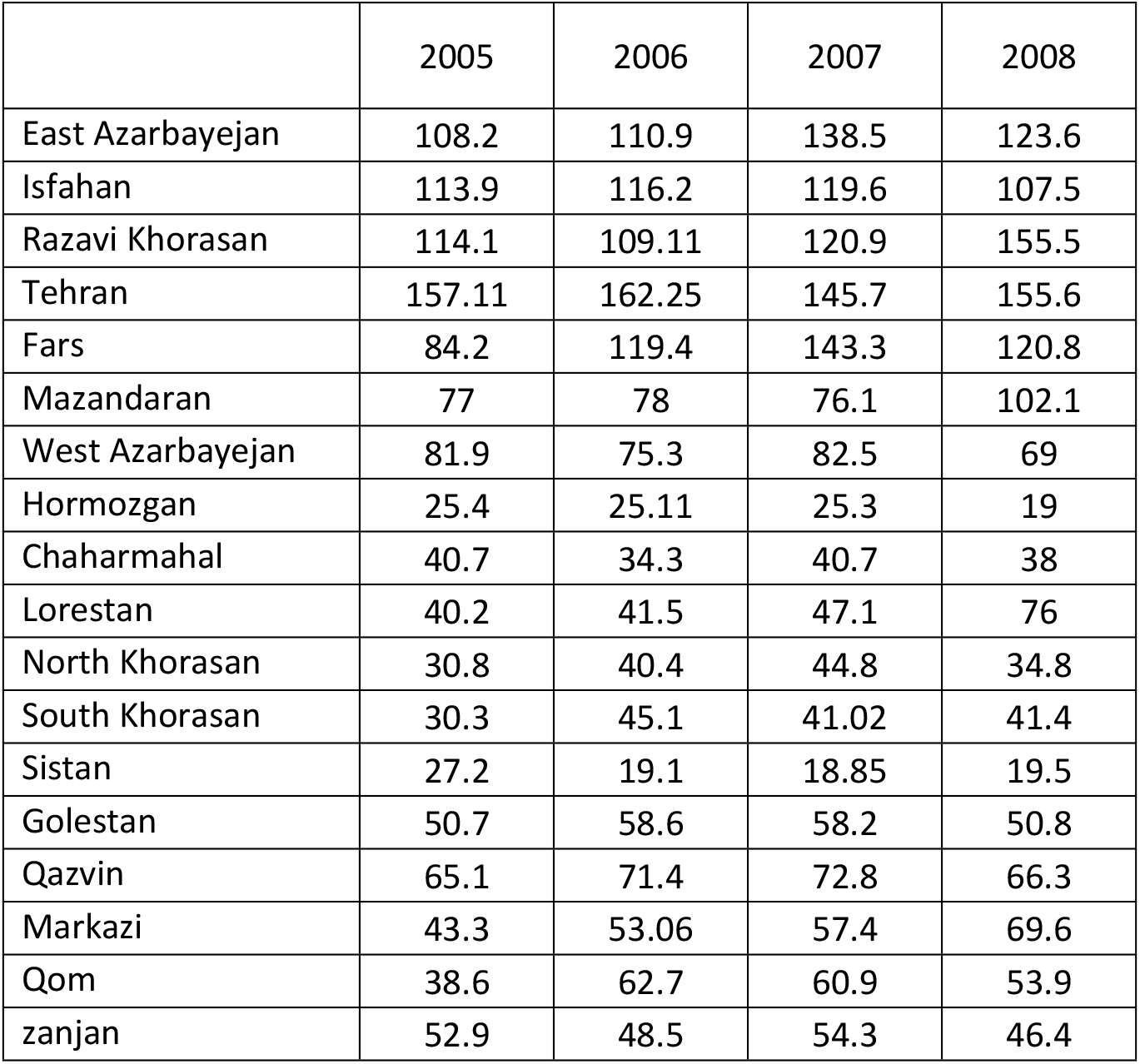
Expected coverage of cancer cases in provinces of Iran (2005-2008).

**Table 2.** Bayesian estimated from misclassification rate between provinces (2005-2008).

|  |  | Estimated misclassification rate |  |  |  |
| --- | --- | --- | --- | --- | --- |
|  |  | 2005 | 2006 | 2007 | 2008 |
| East Azarbayejan | West Azarbayejan | 0.19 | 0.19 | 0.3 | 0.43 |
| Fars | Hormozgan | 0.45 | 0.61 | 0.58 | 0.58 |
| Isfahan | Chaharmahal | 0.44 | 0.43 | 0.38 | 0.47 |
| Isfahan | Lorestan | 0.52 | 0.51 | 0.51 | 0.32 |
| Razavi Khorasan | North Khorasan | 0.57 | 0.66 | 0.53 | 0.55 |
| Razavi Khorasan | South Khorasan | 0.6 | 0.3 | 0.56 | 0.55 |
| Razavi Khorasan | Sistan |  | 0.73 | 0.74 | 0.76 |
| Mazandaran | Golestan | 0.44 | 0.53 | 0.34 | 0.43 |
| Tehran | Qazvin | 0.33 | 0.36 | 0.36 | 0.43 |
| Tehran | Markazi | 0.49 | 0.44 | 0.41 | 0.45 |
| Tehran | Qom | 0.48 | 0.41 | 0.33 | 0.45 |
| Tehran | zanjan | 0.45 | 0.48 | 0.35 | 0.58 |

For example by using the Bayesian method, misclassification rate was estimated 58% between Fars and Hormozgan in 2008. So, after Bayesian correction, ASR and number of cancer incidence decrease for Fars province and increase for Hormozgan province. ASR and number of cancer incidence, before and after Bayesian correction from 2005 to 2008 are reported in tables 3 and 4.

**Table 3.** Age standardized rate of colorectal cancer incidence before and after Bayesian correction in Iranian provinces 2005-2008.

|  | before Bayesian correction |  |  |  | after Bayesian correction |  |  |  |
| --- | --- | --- | --- | --- | --- | --- | --- | --- |
|  | 2005 | 2006 | 2007 | 2008 | 2005 | 2006 | 2007 | 2008 |
| East Azarbayejan | 3.52 | 3.26 | 10.68 | 14.15 | 2.51 | 1.95 | 8.60 | 11.15 |
| Isfahan | 8.19 | 9.26 | 9.05 | 9.54 | 4.74 | 5.00 | 5.78 | 4.14 |
| Razavi Khorasan | 6.10 | 9.44 | 11.04 | 12.96 | 3.53 | 5.32 | 7.04 | 9.65 |
| Tehran | 9.26 | 10.78 | 7.56 | 11.60 | 7.30 | 9.21 | 3.55 | 6.64 |
| Fars | 5.10 | 6.65 | 9.96 | 18.71 | 3.30 | 5.28 | 8.39 | 16.59 |
| Mazandaran | 9.03 | 10.08 | 10.36 | 12.76 | 6.30 | 6.20 | 13.32 | 9.21 |
| West Azarbayejan | 5.31 | 6.46 | 7.59 | 6.30 | 6.54 | 8.09 | 10.35 | 10.23 |
| Hormozgan | 3.46 | 3.00 | 4.19 | 4.99 | 9.59 | 10.29 | 13.80 | 20.23 |
| Chaharmahal | 6.00 | 7.83 | 10.15 | 7.55 | 12.49 | 17.65 | 19.63 | 16.88 |
| Lorestan | 3.96 | 4.52 | 5.45 | 9.52 | 9.08 | 10.07 | 11.35 | 13.53 |
| North Khorasan | 3.00 | 1.70 | 3.20 | 5.02 | 8.55 | 4.48 | 6.99 | 12.94 |
| South Khorasan | 2.48 | 2.89 | 2.88 | 4.40 | 7.39 | 4.81 | 6.81 | 10.24 |
| Sistan | 1.79 | 2.10 | 1.75 | 1.86 | 6.20 | 10.13 | 8.62 | 9.09 |
| Golestan | 5.44 | 7.69 | 9.12 | 7.73 | 10.16 | 14.65 | 14.45 | 14.27 |
| Qazvin | 7.92 | 6.66 | 5.80 | 9.23 | 11.93 | 10.02 | 8.67 | 15.22 |
| Markazi | 4.88 | 6.19 | 6.16 | 7.58 | 10.40 | 11.32 | 10.56 | 12.48 |
| Qom | 5.66 | 6.06 | 9.06 | 9.49 | 12.70 | 10.02 | 13.97 | 17.42 |
| zanjan | 5.43 | 5.38 | 9.21 | 5.23 | 10.05 | 10.70 | 15.15 | 11.77 |

**Table 4.** Number of colorectal cancer incidence and the percent of change before and after Bayesian correction in Iranian provinces 2005-2008.

|  | before Bayesian correction |  |  |  | after Bayesian correction |  |  |  |
| --- | --- | --- | --- | --- | --- | --- | --- | --- |
|  | 2005 | 2006 | 2007 | 2008 | 2005 | 2006 | 2007 | 2008 |
| East Azarbayejan | 95 | 85 | 293 | 379 | 68 | 51 | 236 | 299 |
| Isfahan | 268 | 312 | 279 | 302 | 155 | 168 | 178 | 131 |
| Razavi Khorasan | 307 | 396 | 377 | 432 | 178 | 223 | 241 | 322 |
| Tehran | 819 | 1034 | 315 | 481 | 646 | 883 | 148 | 276 |
| Fars | 166 | 212 | 951 | 1870 | 108 | 132 | 801 | 1658 |
| Mazandaran | 192 | 214 | 217 | 274 | 134 | 132 | 279 | 198 |
| West Azarbayejan | 118 | 135 | 157 | 129 | 145 | 169 | 214 | 209 |
| Hormozgan | 33 | 33 | 44 | 56 | 91 | 113 | 145 | 227 |
| Chaharmahal | 41 | 46 | 65 | 50 | 85 | 104 | 126 | 112 |
| Lorestan | 53 | 70 | 70 | 115 | 122 | 156 | 146 | 163 |
| North Khorasan | 19 | 10 | 18 | 31 | 54 | 26 | 39 | 80 |
| South Khorasan | 9 | 11 | 12 | 18 | 27 | 18 | 28 | 42 |
| Sistan | 31 | 39 | 33 | 34 | 107 | 188 | 163 | 167 |
| Golestan | 67 | 91 | 106 | 90 | 125 | 173 | 168 | 166 |
| Qazvin | 59 | 57 | 48 | 75 | 89 | 86 | 72 | 124 |
| Markazi | 49 | 65 | 63 | 76 | 104 | 119 | 108 | 125 |
| Qom | 44 | 47 | 72 | 74 | 99 | 78 | 111 | 136 |
| zanjan | 39 | 38 | 66 | 42 | 72 | 76 | 109 | 95 |

## Discussion

It is obvious, neighboring provinces due to having the same food habitation, lifestyle and locating in the same climate, have the same health outcomes [13]. But sometimes when analyzing registered data, it is observed that the neighboring provinces not only do not have the same outcomes but also have inconsistent. This situation implies that there is misclassification in registered data. This problem is a notable matter in medicine issues which may results to deflection in health programing and health resources allocating. Such deflection would make irrecoverable damage in national scale. The aim of the present study was to help to reduce misclassification error in registered colorectal cancer data in Iran. In Iran, facility in accessing health resources in welfare provinces at first and secondly lack of health facilities in their neighboring provinces are elements creating misclassification error. Fortunately, some studies have been conducted in Iran in order to eliminate the misclassification errors for mortality and morbidity registered cancer data in the case of Liver [29], Gastric [22], and colorectal cancers [23]. Since the above studies had re-estimated data and also had produced valid data, employing their results may be more reliable. According to the result of our research, there was a non-ignorable estimated misclassification rate among adjacent provinces. The highest estimated misclassification parameter, was belong to North Khorasan, Hormozgan, and Sistan which are in east and south of Iran. So the real rates of CRC in those provinces are higher than the rates that are reported by cancer registry system.

On the contrary, in studies that are used cancer registry data ignoring the existence of misclassification error, it is reported that, the highest incidence rates of CRC in Iran were found in the central, northern, and western provinces; and the southwest provinces of Iran had the lowest incidence rates of CRC in the country! [2]. So, ignoring the misclassification error in registry data, leads to a wrong image of distribution of CRC incidence across the country. Expected cancer coverage revealed that from 30 provinces, 18 provinces need to misclassification correction. These provinces are those which are different in economic situation and there are some points in them which are welfare and probably patients for better health care, refers to those welfare places, so they have more referring people than their capacity. On the other hand, some provinces due to less facility, have less referring patients. Table 2 is indicating how the data of some provinces are registered in their adjacent locations. For example, some neighboring Tehranian patients such as Qom’s patients, were referred to Tehran.

Identifying the exact distribution of disease in different areas is a good manner for finding the geographic pattern of disease and causations, assessing the influencing factors on disease incidence [30,31], and quantifying the potentials for disease control and prevention [32,33]. But usually spatial analysis is used for this purpose which is based on registered data while existence of misclassification is often ignored. In spatial analysis, the morbidity or mortality rates for each province are combined with locality’s information for the same province and so the result may lead to an integrated geographical map. This type of maps is helpful in comparison between different provinces in aspect of rate of disease incidence or probable risk factors [34]. For accessing such a goal, we have prepared geographical map for evaluating incidence distribution of colorectal cancer registered data in before and after misclassification correction in Fig 2. Fig 2 revealed that after correction the southern provinces have high incidence rate, while in the previous studies which had been not regarded misclassification, southern provinces had low incidence rate [35].

**Fig 2.**
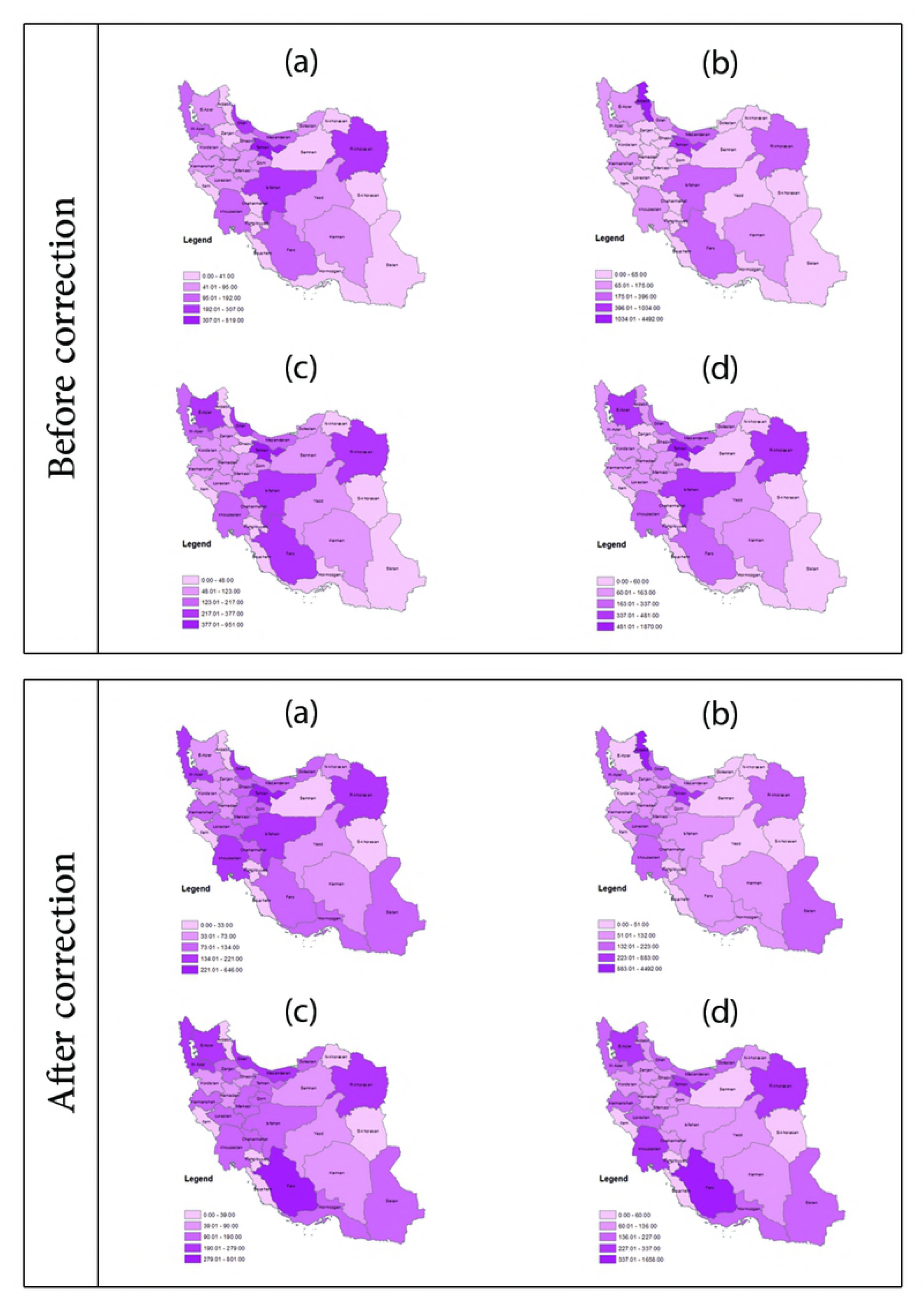
Distribution of Colorectal cancer incidence in Iran before and after misclassification error correction since 2005 to 2009. (a) 2005, (b) 2006, (c) 2007, (d) 2008.

The maps of present study also revealed that a considerable changes happened in some provinces respect to before correction status. Thus major differences in the incidence of CRC, while it is expected that the incidence of cancer be alike in adjacent provinces, can be justified by existence of misclassification error in registering permanent address of patients who are diagnosed in neighboring facilitate provinces. It leads to overestimation of CRC rate in some provinces and underestimation of its rate in some neighboring provinces.

For future researches, to recognizing high risk spatial clusters, using our colorectal cancer valid data, is suggested.

In conclusion, proper planning for cancer control and prevention, and allocating healthcare facilities to different areas, requires an increase in the quality and accuracy of registering system in different provinces, and correcting the existed deficiencies especially misclassification error in registering patient’s permanent residence. It is in need of enhancing hardware and software resources, training more of educated staff in different sectors of cancer registry program, and implementing the opinions of expert researchers in medicine, biostatistics, and epidemiology [36]. In the absence of valid data, Bayesian method can be adopted as a fast and cost effective method to correct the regional misclassification error.

## Acknowledgement

Research reported in this publication was supported by Elite Researcher Grant Committee under award number 958696 from the National Institutes for Medical Research Development (NIMAD), Tehran, Iran

## References

1. Ferlay J, Soerjomataram I, Ervik M, Dikshit R, Eser S, Mathers C, et all. Cancer incidence and mortality worldwide: sources, methods and major patterns in GLOBOCAN 2012. Int J Cancer. 2015; 136(5):E359–86. https://doi.org/10.1002/ijc.29210 PMID: 25220842

2. Khosravi Shadmani F, Ayubi E, Khazaei S, Sani M, Mansouri Hanis S, Khazaei S, et all. Geographic distribution of the incidence of colorectal cancer in Iran: a population-based study. Epidemiology and health. 2017; 39. http://doi.org/10.4178/epih.e2017020. PMID: 28774167

3. Mohagheghi MA, Mosavi-Jarrahi A, Malekzadeh R, Parkin M. Cancer incidence in Tehran metropolis: the first report from the Tehran Population-based Cancer Registry, 1998-2001. Arch Iran Med. 2009; 12(1):15–23. PMID: 19111024

4. Johnson CM, Wei C, Ensor JE, Smolenski DJ, Amos CI, Levin B, et al. Meta-analyses of colorectal cancer risk factors. Cancer Causes Control. 2013; 24(6):1207–22. http://doi.org/10.1007/s10552-013-0201-5. PMID: 23563998

5. Larsson SC, Orsini N, Wolk A. Diabetes mellitus and risk of colorectal cancer: a meta-analysis. J Natl Cancer Inst. 2005; 97(22):1679–87. http://doi.org/10.1093/jnci/dji375. PMID: 16288121

6. Arnold M, Karim-Kos HE, Coebergh JW, Byrnes G, Antilla A, Ferlay J, et all. Recent trends in incidence of five common cancers in 26 European countries since 1988: Analysis of the European Cancer Observatory. Eur J Cancer. 2015; 51(9):1164–87. http://doi.org/10.1016/j.ejca.2013.09.002. PMID: 24120180

7. Mathers C, Fat DM, Boerma JT. The global burden of disease: 2004 update. World Health Organization;2008.

8. Parkin DM. The evolution of the population-based cancer registry. Nat Rev Cancer. 2006; 6(8):603–12. http://doi.org/10.1038/nrc1948. PMID: 16862191

9. Das A. Cancer registry databases: an overview of techniques of statistical analysis and impact on cancer epidemiology. Methods Mol Biol. 2009; 471:31–49. http://doi.org/10.1007/978-1-59745-416-2_2. PMID: 19109773

10. Pourhoseingholi MA, Vahedi M, Moghimi-Dehkordi B, Pourhoseingholi A, Ghafarnejad F, Maserat E, et all. Burden of hospitalization for gastrointestinal tract cancer patients-Results from a cross-sectional study in Tehran. Asian Pac J Cancer Prev. 2009; 10(1):107–10. PMID: 19469635

11. Yavari P, Sadrolhefazi B, Mohagheghi MA, Madani H, Mosavizadeh A, Nahvijou A, Mehrabi Y, Pourhseingholi M. An epidemiological analysis of cancer data in an Iranian hospital during the last three decades. Asian Pac J Cancer Prev. 2008; 9(1): 145–50. PMID: 18439094

12. Mohagheghi MA, Mosavi-Jarrahi A. Review of cancer registration and cancer data in Iran, a historical prospect. Asian Pac J Cancer Prev. 2010; 11(4):1155–7. PMID: 21133641

13. Islamic Repulic of Iran. Ministry of Health and Medical Education. Center for Disease Control & Prevention. Noncommunicable Deputy. Cancer Office. Iranian Annual of National Cancer Registration Report; 2009.

14. Lyles RH. A note on estimating crude odds ratios in case–control studies with differentially misclassified exposure. Biometrics. 2002; 58(4):1034–36. PMID: 12495160

15. Corbin M. Bayesian methods to address multiple comparisons and misclassification bias in studies of occupational and environmental risks of cancer: a thesis by publications presented in partial fulfilment of the requirements for the degree of Doctor of Philosophy in Public Health. Massey University, Wellington, New Zealand;2013.

16. Whittemore AS, Gong G. Poisson regression with misclassified counts: application to cervical cancer mortality rates. J R Stat Soc Ser C Appl Stat. 1991; 40(1):81–93. PMID: 12157994

17. Sposto R, Preston DL, Shimizu Y, Mabuchi K. The effect of diagnostic misclassification on non-cancer and cancer mortality dose response in A-bomb survivors. Biometrics. 1992; 48(2):605–17. PMID: 1637983

18. McInturff P, Johnson WO, Cowling D, Gardner IA. Modelling risk when binary outcomes are subject to error. Stat Med. 2004; 23(7):1095–109. http://doi.org/10.1002/sim.1656. PMID: 15057880

19. Stamey JD, Young DM, Seaman JW. A Bayesian approach to adjust for diagnostic misclassification between two mortality causes in Poisson regression. Stat Med. 2008; 27(13):2440–52. http://doi.org/10.1002/sim.3134. PMID: 17979218

20. Aghajani H, Eatemad K, Goya M, Ramezani R, Modirian MN, Nadali F. Iranian annual of national cancer registration report 2008-2009. Center for Disease Control;2011.

21. Ahmad OB, Boschi-Pinto C, Lopez AD, Murray CJ, Lozano R, Inoue M. Age standardization of rates: a new WHO standard. Global Programme on Evidence for Health Policy Discussion Paper Series: no. 31. World Health Organization;2001.

22. Hajizadeh N, Pourhoseingholi MA, Baghestani AR, Abadi A, Zali MR. Bayesian adjustment for over-estimation and under-estimation of gastric cancer incidence across Iranian provinces. World J Gastrointest Oncol. 2017; 9(2):87–93. http://doi.org/10.4251/wjgo.v9.i2.87. PMID: 28255430

23. Pourhoseingholi MA, Faghihzadeh S, Hajizadeh E, Abadi A, Zali MR. Bayesian estimation of colorectal cancer mortality in the presence of misclassification in Iran. Asian Pac J Cancer Prev. 2009; 10(4):691–4. PMID: 19827896

24. Hajizadeh N, Baghestani AR, Pourhoseingholi MA, Ashtari S, Fazeli Z, Vahedi M, Zali MR. Trend of hepatocellular carcinoma incidence after Bayesian correction for misclassified data in Iranian provinces. World J Hepatol. 2017; 9(15):704–10. http://doi.org/10.4254/wjh.v9.i15.704. PMID: 28596818

25. Pourhoseingholi MA. Bayesian adjustment for misclassification in cancer registry data. Translational Gastrointestinal Cancer. 2014; 3(4):144–8. http://doi.org/10.3978/j.issn.2224-4778.2014.08.08.

26. Paulino CD, Soares P, Neuhaus J. Binomial regression with misclassification. Biometrics. 2003; 59(3):670–5. PMID: 14601768

27. Paulino CD, Silva G, Achcar JA. Bayesian analysis of correlated misclassified binary data. Computational statistics & data analysis. 2005; 49(4):1120–31. https://doi.org/10.1016/j.csda.2004.07.004.

28. Liu Y, Johnson WO, Gold EB, Lasley BL. Bayesian analysis of risk factors for anovulation. Stat Med. 2004; 23(12):1901–19. http://doi.org/10.1002/sim.1773. PMID: 15195323

29. Hajizadeh N, Baghestani AR, Pourhoseingholi MA, Najafimehr H, Fazeli Z, Bosani L. Bayesian correction model for over-estimation and under-estimation of liver cancer incidence in Iranian neighboring provinces. Gastroenterol Hepatol Bed Bench. 2017; 10(Suppl 1):S54–S61. PMID: 29511473

30. Mehrabani D, Tabei SZ, Heydari ST, Shamsina SJ, Shokrpour N, Amini M, et all. Cancer occurrence in Fars Province, Southern Iran. Iranian Red Crescent Medical Journal. 2008; 10(4):314–22.

31. Kamangar F, Dores GM, Anderson WF. Patterns of cancer incidence, mortality, and prevalence across five continents: defining priorities to reduce cancer disparities in different geographic regions of the world. J Clin Oncol. 2006; 24(14):2137–50. http://doi.org/10.1200/JCO.2005.05.2308. PMID: 16682732

32. Pandey S, Mishra M, Chandrawati C. Human papillomavirus screening in north Indian women. Asian Pac J Cancer Prev. 2012; 13(6):2643–6. PMID: 22938435

33. Zou L, Bao YP, Li N, Dai M, Ma CP, Zhang YZ, et all. Life-style and genital human papillomavirus in a cross-sectional survey in Shanxi Province, China. Asian Pac J Cancer Prev. 2011; 12(3):781–6. PMID: 21627383

34. Zayeri F, Kavousi A, Najafimehr H. Spatial analysis of Relative Risks for skin cancer morbidity and mortality in Iran, 2008-2010. Asian Pac J Cancer Prev. 2015; 16(13):5225–31. PMID: 26225657

35. Mahaki B, Mehrabi Y, Kavousi A, Akbari ME, Waldhoer T, Schmid VJ, et all. Multivariate Disease Mapping of Seven Prevalent Cancers in Iran using a Shared Component Model. Asian Pacific J Cancer Prev. 2011; 12(9):2353–8. PMID: 22296383

36. Teppo L, Pukkala E, Lehtonen M. Data quality and quality control of a population-based cancer registry. experience in Finland. Acta Oncol. 1994; 33(4):365–9. PMID: 8018367

